# A Highly Contiguous Reference Genome for *Scalesia gordilloi* (Asteraceae), a Critically Endangered Plant Endemic to the Galapagos Islands

**DOI:** 10.64898/2026.06.25.734018

**Authors:** Gabriela Pozo, Gonzalo Rivas-Torres, Emilio Vélez-Darquea, Doménica Barragán-Orbe, Maria de Lourdes Torres

## Abstract

*Scalesia gordilloi* is a critically endangered species endemic to San Cristóbal Island in the Galapagos archipelago and represents one of the most unique and vulnerable lineages within the adaptive radiation of the genus *Scalesia*. Despite its evolutionary distinctiveness and conservation importance, no genomic resources have been available for this species. Here, we present the first high-quality reference genome of *S. gordilloi*, generated using Oxford Nanopore long-read sequencing. Across three PromethION R10.4.1 flow cells, we obtained 80.5 Gb of long reads (∼25× coverage), which enabled a highly contiguous 3.61 Gb assembly composed of only 549 contigs and an N50 of 106.6 Mb. BUSCO completeness reached 98.6%, with assembly metrics comparable to other high-quality Asteraceae genomes. Repeat annotation revealed that 76.2% of the genome is composed of interspersed elements, dominated by LTR retrotransposons. Structural annotation resulted in 47,913 high-confidence protein-coding genes, consistent with expectations for large, repetitive Asteraceae genomes. This genome provides a critical foundation for conservation genomics, enabling assessments of genetic diversity, inbreeding, and adaptive potential in the species. It further establishes a framework for comparative genomics across the *Scalesia* radiation and supports future efforts to protect and restore one of the most threatened plant lineages of the Galapagos Islands.

## 1. Background

The *Scalesia* genus (Family Asteraceae, Order Asterales) is an endemic group from the Galapagos archipelago [1]. It is composed of 12 shrubby species and 3 arboreal species that are distributed allopatrically (Itow, 1995). Like Darwin’s finches, species of the genus *Scalesia* are apparently the result of an accelerated process of diversification and adaptation, known as adaptive radiation [2]. This process is common in oceanic islands and has been an important system to study evolutionary mechanisms, since many islands are geologically new [2]. Due to their allopatric distribution, species grow in populations separated from each other, and some are even limited to specific islands, such as *Scalesia gordilloi* [1], which is confined to the southeastern part of San Cristóbal Island (Fig. 1).

**Figure 1.**
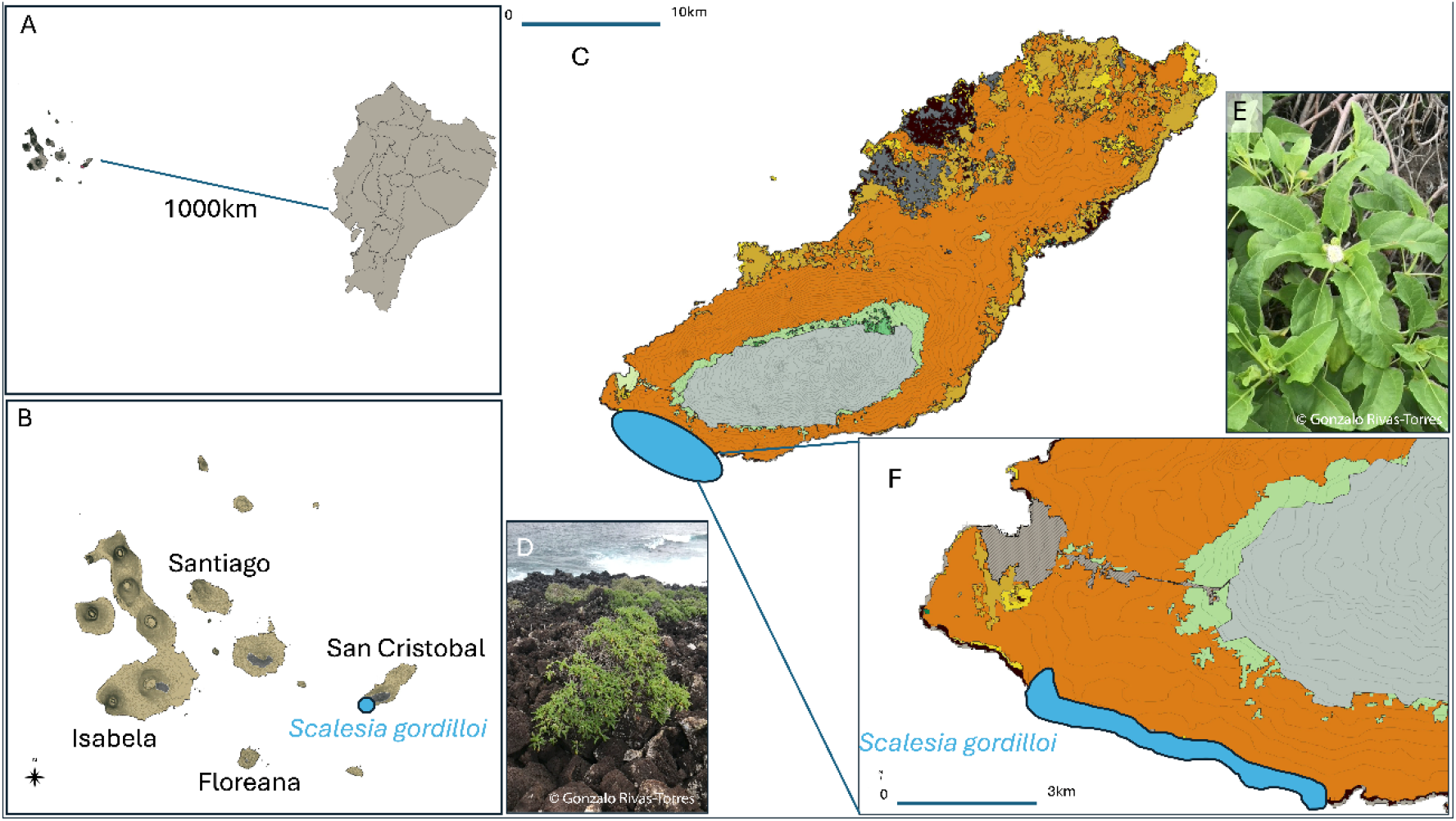
Geographic distribution and habitat of *Scalesia gordilloi* on San Cristóbal Island. (A) Location of the Galapagos Archipelago in relation to mainland Ecuador. (B) Location of San Cristóbal Island in the eastern region of the Galapagos, with Santiago, Isabela, and Floreana Islands shown for reference. (C) Detailed map of San Cristóbal indicating the extent of the Galapagos Dry Forest (in orange) and the approximate distribution of *S. gordilloi* (blue circle). (D) Photograph of an individual of *S. gordilloi* and its native habitat. (E) Close-up image of *S. gordilloi* showing leaf morphology and inflorescence. (F) Detailed map of the known distribution of *S. gordilloi* on San Cristóbal Island.

*Scalesia gordilloi* (Asteraceae/Compositae) was formally described in 1986 [3]. It is characterized as a single-stemmed shrub reaching approximately 1.2–1.5 m maximum in height, although individuals may occasionally exhibit a more prostrate growth form. Its leaves are entire, with petioles measuring approximately 2–3 cm and leaf blades approximately 5–7 cm [3]. The leaves bear glandular structures and, when crushed, release a somewhat pleasant aroma, a characteristic maintained in most of the *Scalesia* species [4].

The species possesses capitulate inflorescences, with individuals being partially heterogamous. The florets are white, and the heads measure approximately 8–10 mm in length (Fig. 1D) [3]. The achenes are about 3 mm in maximum length and 1 mm at their widest point, and dispersal occurs primarily by wind.

As mentioned, *S. gordilloi* is a species with a highly restricted distribution, as it has been recorded exclusively on San Cristóbal Island, making it endemic to this island. Within San Cristóbal, *S. gordilloi* occurs in just over 400 hectares in the southwestern portion of the island (G. Rivas-Torres, unpublished data), one of the easternmost and geologically oldest of the Galapagos archipelago (Fig. 1C). The species occurs as scattered individuals (approximately every 20–40 m apart) within this range, and is primarily distributed within the Galapagos Deciduous Forest, a dry ecosystem that occupies the lower-elevation perimeter of the islands and is characterized by comparatively lower precipitation, lower relative humidity, and reduced organic soil content than the more humid highland ecosystems [5].

Although *S. gordilloi* is one of four *Scalesia* species present on San Cristóbal, its distribution does not appear to overlap with that of any other *Scalesia* species. *S. gordilloi* is restricted to the southern coast of San Cristóbal, where it occurs in low-elevation arid coastal habitats (Hamann & Wium-Andersen, 1986). In these coastal habitats, it is regularly exposed to sea spray from breaking waves, which often results in the presence of salt crystals on the leaf surfaces. Current population estimates suggest that the species consists of only a few hundred individuals, primarily mature adults (G Rivas-Torres, personal observation). The species is currently listed as Critically Endangered on the IUCN Red List [6]. *S. gordilloi* represents one of the more late-diverging lineages within the genus and is closely related to the more evolutionarily young *Scalesia* species, including *S. divisa* and *S. incisa*, which also occur on San Cristóbal but are found further to the north on the island (Fernández-Mazuecos et al. 2020).

The Galapagos Islands harbor a unique biodiversity, being the habitat of iconic species, such as giant tortoises or Darwin’s finches [7]. However, the introduction of non-native species represents one of the greatest threats to local populations [8]. In the case of the genus *Scalesia*, which represents one of the most emblematic examples of adaptive radiation in plants in the archipelago, several species are currently threatened or critically endangered, including *S. gordilloi*. Despite their ecological and evolutionary importance, genomic resources for this group remain scarce. To date, *S. atractyloides*, one of the basal lineages of the group, appears to be the only species with a high-quality reference genome available [9], highlighting an important gap in genomic knowledge for the genus.

To help address these limitations, several initiatives have emerged aiming to expand genomic resources for biodiversity, particularly in understudied regions. One such effort is ORG.one, which focuses on generating de novo genome assemblies for endangered and critically endangered species using Oxford Nanopore sequencing technologies. These initiatives facilitate the rapid production of genomic data and promote open access to these resources, enabling their use in conservation, evolutionary, and ecological studies [10,11].

Obtaining a high-quality reference genome is essential for enabling downstream analyses, including the identification of genetic variation, comparative genomics, and phylogenetic inference [12]. Such resources are particularly valuable for species with small and fragmented populations, as they provide critical insights into genetic diversity, population structure, and adaptive potential, all of which are key components for informed conservation strategies.

To address this gap, we generated a comprehensive genomic dataset for *S. gordilloi* using Oxford Nanopore long-read sequencing as part of the ORG.one initiative. This study provides not only a high-quality reference genome assembly, but also associated raw sequencing data and genome annotation resources. A detailed description of these datasets, including their generation, quality, and potential applications, is provided in the following section.

## 2. Data Description

This study generated a comprehensive genomic dataset for *Scalesia gordilloi*, including raw long-read sequencing data, a de novo genome assembly, and structural genome annotation. The assembled genome spans 3.61 Gb and is composed of 549 contigs, with a contig N50 of 106.6 Mb, indicating a highly contiguous assembly. Genome completeness was assessed using BUSCO, recovering 98.6% of conserved single-copy orthologs. Repetitive element annotation revealed that approximately 76.2% of the genome consists of interspersed repeats, primarily long terminal repeat (LTR) retrotransposons.

Structural annotation resulted in 47,913 high-confidence protein-coding genes following filtering steps to remove short and transposable element–associated predictions. The resulting gene models include exon–intron structures consistent with other large Asteraceae genomes.

These datasets provide a valuable resource for multiple downstream applications, including comparative genomics, studies of genome evolution within the Asteraceae, and conservation genomics of endangered island species. In particular, the availability of a high-quality reference genome enables future analyses of genetic diversity, population structure, and adaptive variation in *S. gordilloi* and related taxa.

All datasets associated with this study, including raw sequencing reads, genome assembly, and annotation files, are available through public repositories (see Data Availability section), ensuring accessibility and reuse by the research community.

## 3. Analyses

### 3.1 Sequencing output and data quality

To assess sequencing output and dataset quality, we generated long-read data using R10.4.1 flow cells on the Oxford Nanopore PromethION 2 Solo platform, obtaining a total of 80.5 Gb of raw sequence data across 8.4 million reads. The dataset showed a mean read length of 9.6 kb (median 7.2 kb), a read length N50 of 15.0 kb, and a mean read quality of 12.9 (median 15.2). Based on the estimated genome size (∼3.2 Gb), the dataset represents approximately 25× long-read coverage, providing sufficient depth to support a robust de novo assembly.

### 3.2 Genome assembly and assembly completeness

To generate a high-quality reference genome for *Scalesia gordilloi*, we performed de novo assembly using Hifiasm, which yielded a total genome size of 3.61 Gb, distributed across 549 contigs (Table 1, Figure 2). The longest contig measured 143.7 Mb, with an N50 of 106.6 Mb and L50 of 16, indicating a highly contiguous assembly. Comparison with the *S. atractyloides* genome showed that 82.2% of the *S. gordilloi* assembly aligned to the reference, reflecting strong genomic conservation within the genus. To evaluate assembly completeness, BUSCO analysis recovered 98.6%), with only 0.4% of genes missing,consistent with a near-complete representation of the *S. gordilloi* genome (Table 1, Figure 2).

**Table 1.** Genome assembly and BUSCO statistics for Scalesia gordilloi.

| **Statistic** | **Assembly** |
| --- | --- |
| **Contig number** | 549 |
| **Longest contig** | 143.7 Mb |
| **Total Length** | 3.61 Gb |
| **N50** | 106.6 Mb |
| **L50** | 16 |
| **Genome fraction\*** | 82.2 % |
| **Complete BUSCOs\*\*** | 98.6% |
| **Single-copy** | 12% |
| **Duplicated** | 86.7% |
| **Fragmented BUSCOs** | 0.9% |
| **Missing BUSCOs** | 0.4% |

\*Percentage of genome that aligns to reference *Scalesia atractyloides*
\*\*Database eudicots\_odb10 with 2,805 orthologous genes

**Figure 2.**
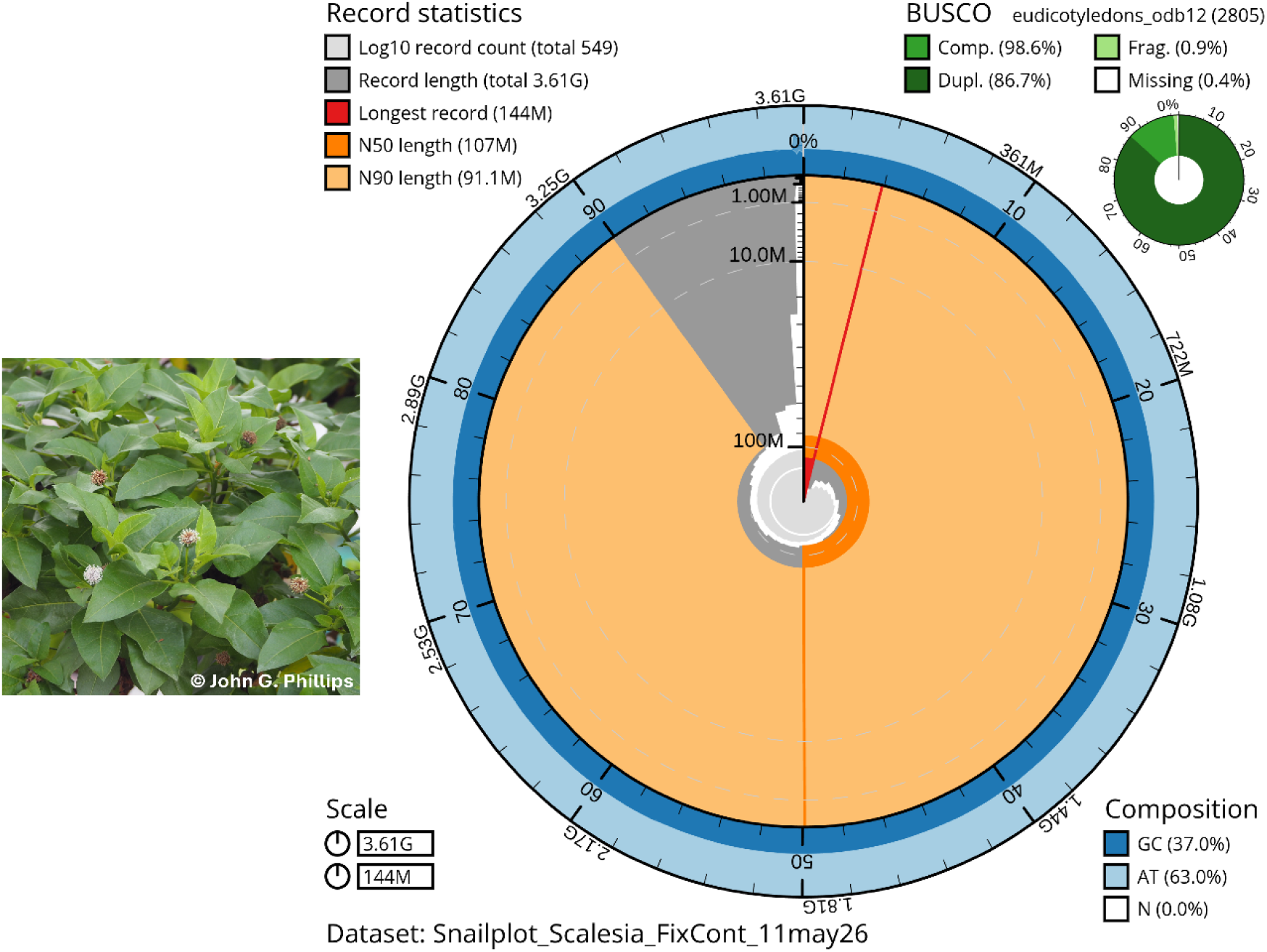
Snailplot summary statistics of the *Scalesia gordilloi* reference genome assembly. Summary shows a total assembly size of 3.61 Gb distributed across 549 contigs. The longest scaffold is 144 Mb, with an N50 of 107 Mb and N90 of 91.1 Mb, indicating a highly contiguous assembly. BUSCO analysis using the eudicotyledons dataset recovered 98.6% complete genes (86.7% duplicated), with 0.9% fragmented and 0.4% missing. GC content is 37.0% and AT content is 63.0%.

### 3.3 Repeat annotation

To characterize genome composition, we identified repetitive elements, which accounted for 76.2% of the *S. gordilloi* genome (Table 2). Among these, retroelements represented the dominant class, representing 39.6% of the genome, followed by unclassified repeats (34.9%) and DNA transposons (1.7%). Long terminal repeat (LTR) elements were particularly abundant, especially Gypsy/DIRS1 and Ty1/Copia families, comprising 23.8% and 11.0%, respectively.

**Table 2.**
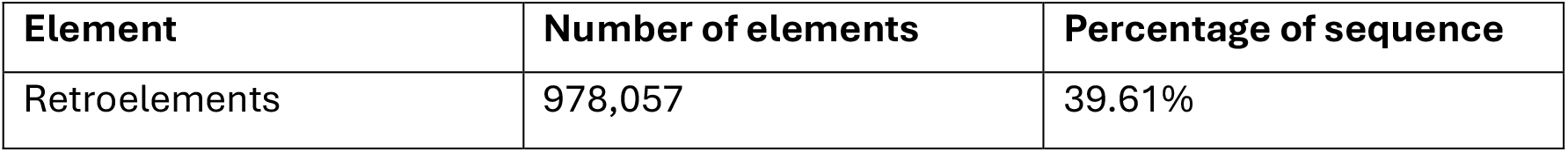
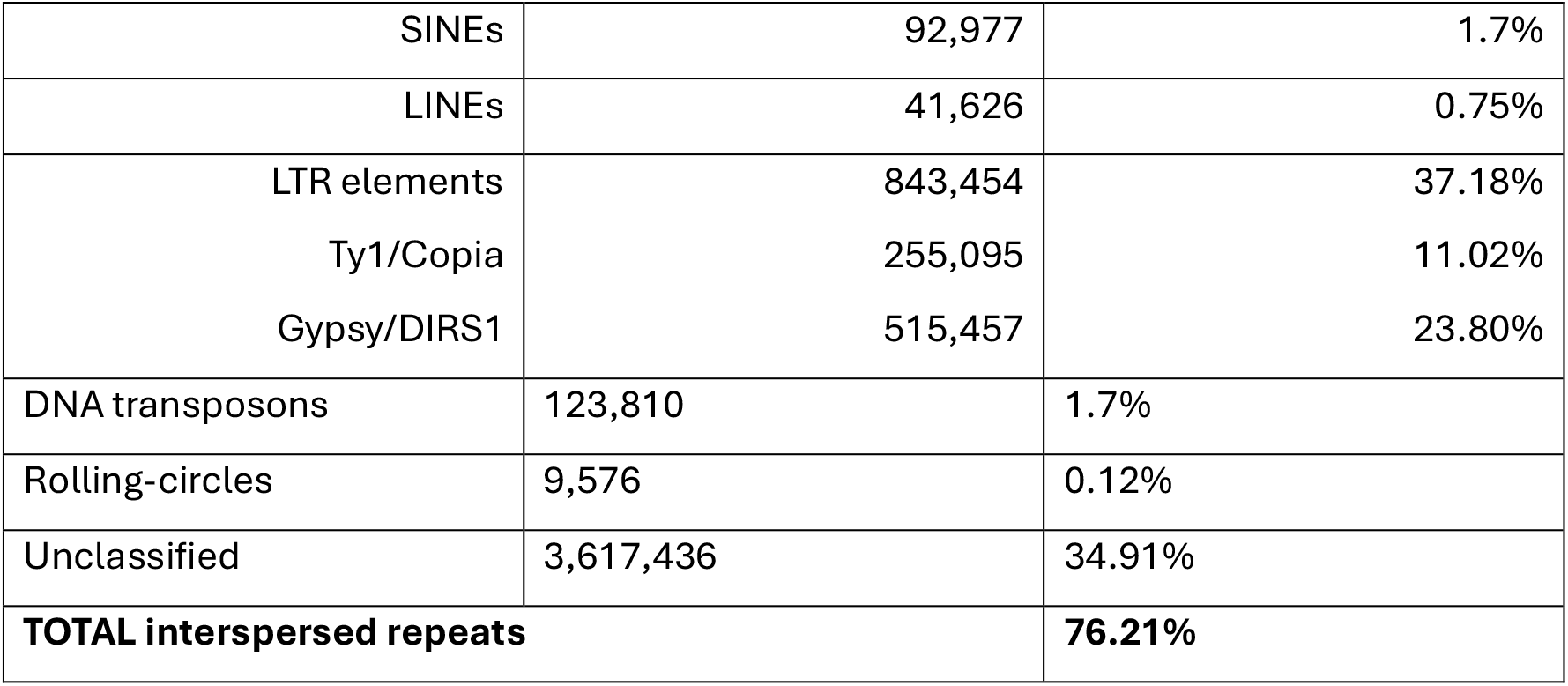
Repetitive elements in the Scalesia gordilloi reference genome.

| Element | Number of elements | Percentage of sequence |
| --- | --- | --- |
| Retroelements | 978,057 | 39.61% |
| SINEs | 92,977 | 1.7% |
| LINEs | 41,626 | 0.75% |
| LTR elements | 843,454 | 37.18% |
| Ty1/Copia | 255,095 | 11.02% |
| Gypsy/DIRS1 | 515,457 | 23.80% |
| DNA transposons | 123,810 | 1.7% |
| Rolling-circles | 9,576 | 0.12% |
| Unclassified | 3,617,436 | 34.91% |
| **TOTAL interspersed repeats** |  | **76.21%** |

### 3.4 Structural annotation

To characterize gene content, structural annotation of the filtered genome yielded 47,913 protein-coding genes, each represented by a single transcript. Genes contained an average of 5.4 exons and 4.4 introns, with mean exon and intron lengths of 308 bp and 332 bp, respectively (Table 3). Coding sequences averaged 1,674 bp, and gene models ranged from 906 bp to 59.9 kb in total length. A total of 9,342 genes (19.5%) were single-exon models. The final annotation encompassed 80.2 Mb of coding sequences, representing the coding fraction of the assembled genome.

**Table 3.** Structural genome annotation statistics.

| Statistic | Value |
| --- | --- |
| Number of genes | 47,913 |
| Number of exons | 259,933 |
| Number of introns in CDS | 212,020 |
| Mean exons per gene | 5.4 |
| Mean introns per gene | 4.4 |
| Mean gene length (bp) | 3,145 bp |
| Gene size range | 906 – 59,899 bp |
| Mean exon length (bp) | 308 bp |
| Mean intron length (bp) | 332 bp |
| Single-exon genes | 9,342 (19.5%) |
| Mean CDS length | 1,674 bp |
| Longest CDS | 15,231 bp |
| Total CDS length | 80.2Mb |

## 4. Discussion

Oxford Nanopore sequencing produced 80.5 Gb of long-read data across 8.4 million reads, equivalent to ∼25× coverage of the estimated *Scalesia gordilloi* genome. The dataset showed a mean read length of 9.6 kb, a read N50 of 15 kb, and a mean quality score of Q12.9, values consistent with high-yield PromethION R10.4.1 runs and suitable for assembling large, repeat-rich plant genomes [13,14]. Long reads are particularly advantageous for spanning repetitive regions, which dominate Asteraceae genomes and which was later shown to constitute ∼76% of the *S. gordilloi* genome. Overall, these sequencing metrics provided sufficient depth and read quality for generating a high-contiguity de novo assembly.

The final *S. gordilloi* assembly is highly contiguous, with a total size of 3.61 Gb, only 549 contigs, and an N50 of 106.6 Mb. These metrics represent a substantial improvement over earlier drafts and approach the contiguity of the chromosome-level genome of *S. atractyloides* [9]. BUSCO completeness was 98.6%, with only 0.4% missing genes, matching the completeness of other high-quality Asteraceae genomes such as sunflower and lettuce [13,14]. The high proportion of duplicated BUSCOs is consistent with known whole-genome duplication events and TE-driven expansion in the family [9,13–15]. Together, these metrics indicate that the assembly captures both the gene-rich and repeat-dense regions of the genome with high accuracy.

When aligning our assembly to the published genome of closely related *S. atractyloides* (GCA_947069175.1), we obtained an alignment of approximately 82.2%, indicating a high degree of large-scale genomic similarity between these two *Scalesia* species despite their phylogenetic and geographic differences [2]. Consistent with this result, QUAST and BUSCO metrics support the high quality and completeness of the assembly, indicating its suitability for downstream applications, including comparative genomics, repeat evolution analyses, and population-level studies.

The final *S. gordilloi* annotation comprises 47,913 protein-coding genes, a number consistent with expectations for large, highly repetitive genomes within the Asteraceae, many of which contain between 40,000 and 55,000 genes due to extensive transposable-element expansion and historical whole-genome duplications [13,14]. This gene count also closely matches that of *S. atractyloides*, the only previously published chromosome-level genome for the genus. In that species, Cerca et al. (2022) reported 46,375 Iso-Seq– supported genes and 43,093 final annotated, providing a strong benchmark for the genomic architecture of *Scalesia* [9]. Overall, the similarity in gene number between the two *Scalesia* species is consistent with broadly comparable genome organization, although additional genus-wide analyses would be necessary to evaluate whether this reflects shared evolutionary patterns across the genus.

The annotation statistics of *S. gordilloi* are fully consistent with other Asteraceae genomes, which are characterized by high repeat content (>70%) and large gene repertoires. For example, sunflower (*Helianthus annuus*) contains ∼52,000 genes and 74% repeats [13], lettuce (*Lactuca sativa*) contains ∼39,000 genes and 74% repeats [14], and the Hawaiian *Bidens hawaiensis* genome harbors 74% repetitive DNA [15]. The ∼76% repeat content and ∼48k genes recovered in *S. gordilloi* therefore fall well within the expected variation for this family. Additionally, the recovery of well-structured exon–intron architectures and the absence of widespread gene fragmentation further support the robustness of the annotation.

According to this study, *S. gordilloi* and its more distantly related congener *S. atractyloides* share approximately 82.2% of their genome. While direct genome-wide similarity estimates are rarely reported for plants, studies in Asteraceae and other angiosperms indicate that closely related species often exhibit high gene-level similarity (typically 85–95% identity among orthologs) alongside substantial structural variation [13,14,16]. Therefore, this value likely reflects a moderate-to-high level of genomic similarity, although differences in methodology and assembly quality limit direct comparisons. However, despite this genomic similarity, the most recent and comprehensive phylogenetic analyses indicate that although they share a common ancestor, these species do not cluster in closely related clades. *S. atractyloides* is more closely related to *S. stewartii*, and together these two species form a monophyletic clade that is evolutionarily closer to the ancestral and more basal species *S. villosa* [2]. Moreover, *S. atractyloides* and *S. gordilloi* occur on islands that are geographically distant from one another. *S. atractyloides* is distributed in the southwestern region of Santiago Island, which lies approximately 152 km northwest of the *S. gordilloi* population on San Cristóbal. This distance is separated almost entirely by open ocean. However, we did not directly assess large-scale genomic structure; additional analyses such as synteny would be needed to evaluate structural conservation across the genus.

Overall, the *S. gordilloi* genome and gene models constitute a robust and well-supported genomic resource, exhibiting characteristics comparable to those reported for other published Asteraceae genomes. This high-quality resource provides a strong foundation for future studies of comparative genomics, adaptive evolution, and island diversification within this endemic Galapagos lineage.

This study also provides a scalable methodology for generating reference genomes of additional *Scalesia* species. A clear next step is sequencing the close relatives of *S. gordilloi, S. incisa* and *S. divisa*, which are listed as Vulnerable and Critically Endangered, respectively [17,18]. Obtaining high-quality genomes for these species would refine their phylogenetic relationships, improve assessments of genomic diversity and conservation value, and enable identification of genes linked to ecological specialization. Such genomic resources will be critical for informing conservation actions, predicting adaptive potential under climate change, and guiding restoration strategies across the genus.

## 5. Potential Implications

The high-quality reference genome of *Scalesia gordilloi* provides a valuable resource that can be applied beyond the scope of this study. In conservation genomics, this dataset enables future analyses of genetic diversity, inbreeding, adaptive variation, and population structure in one of the most threatened plant species of the Galapagos Islands. Such information will be essential for informing evidence-based conservation strategies, including habitat restoration, population monitoring, ex situ conservation, and potential assisted gene flow approaches. In addition, the availability of a reference genome may support the identification of genetically vulnerable populations and help guide conservation actions aimed at preserving the evolutionary potential and long-term resilience of the species under ongoing environmental change.

More broadly, this genome contributes to comparative genomic studies within the Asteraceae, a family characterized by large and highly repetitive genomes. The availability of this resource will facilitate investigations into genome evolution, transposable element dynamics, and adaptation in island plant radiations. In particular, comparative analyses across the *Scalesia* genus may help elucidate the genomic basis of adaptive radiation in oceanic island systems.

Finally, this study demonstrates a scalable approach for generating high-quality reference genomes for other endemic and endangered plant species using long-read sequencing technologies. The methodologies and datasets presented here can support future genomic efforts in biodiversity-rich but underrepresented regions, promoting the integration of genomics into conservation and evolutionary research.

## 1. Methods

### 6.1 Sample collection

Seven adult individuals of *Scalesia gordilloi* were sampled within the known distribution range of the species. Each sampled plant was located at least 20 meters apart to ensure independence of collections. From each individual, 3–5 young leaves were collected and immediately placed in airtight bags with moist paper to maintain tissue hydration. These bags were then placed in cooled containers in the field to maintain low temperature conditions.

The samples were transported to the Galapagos Science Center, where they were stored at –20 °C. Samples were subsequently transported in sealed cold containers to the Plant Biotechnology Laboratories of USFQ in Quito. All sampling, handling, and DNA extraction procedures were conducted under Research Permit No. MAAE-DBI-CM-2021-0198. No individuals were harmed during sampling, as care was taken to avoid damaging branches or trampling other plants within the population.

### 6.2 DNA Extraction, Library preparation, and Sequencing

High molecular weight DNA was obtained from fresh young leaf tissue of a unique *S. gordilloi* specimen using the Blood & Cell Culture DNA Kit (Qiagen) and a modified CTAB protocol [19] starting with 1 g and 100 mg of tissue, respectively. DNA was eluted in 100 µl, and fragments shorter than 25 kb were removed using the Circulomics Short Read Eliminator Kit (PacBio). Finally, DNA concentration was measured using a Qubit 4 Fluorometer and DNA integrity was evaluated with electrophoresis on a 1.5% agarose gel.

The Ligation Sequencing Kit V14 (SQK-LSK114; Oxford Nanopore Technologies) was used to prepare one sequencing library for each DNA extraction method, using 1000 ng of initial DNA material. The libraries were prepared according to the manufacturer’s protocol for the Ligation Sequencing Kit V14, with a few small modifications, specifically during the DNA repair and end-prep step (15 minutes of incubation at 20°C, 15 minutes at 65°C, and a single ethanol wash), and the adapter ligation and clean-up step (20 minutes of incubation at room temperature and a single wash with long fragment buffer).

Sequencing was carried out on a PromethION 2 Solo using R10.4.1 flow cells across three runs lasting approximately 24, 46 and 66 hours. MinKNOW (v6.5.14) was used for data collection, and high-accuracy basecalling was performed by Dorado (v7.9.8) with a minimum Q score of 7.

### 6.3 Genome Assembly and Analyses

All reads that passed the Q7 threshold were concatenated prior to adapter removal with Porechop v0.2.3_seqan2.1.1 [20]. Read quality and length were evaluated using NanoPlot v1.32.1, and dataset cleanup was performed with NanoFilt v2.3.0 [21] by removing reads shorter than 2,000 bp. The resulting filtered dataset was used for the *de novo* genome assembly of *S. gordilloi* using Hifiasm v0.25.0-r726 [22], executed with ONT-specific settings. Raw reads were subsequently aligned to the draft assembly using minimap2 v2.30- r1287 [23], and two iterative rounds of polishing were performed with Racon v1.4.20 [24]. Finally, duplicated contigs resulting from haplotypes in polyploid genomes were identified and removed with Purge Haplotigs v1.1.3 [25]. Assembly completeness was assessed using BUSCO v5.8.3 [26], and assembly statistics were generated with QUAST v5.0.2 [27].

### 6.4 Genome Annotation and Repetitive Elements Analysis

Structural gene annotation was performed using the soft-masked *S. gordilloi* genome produced with RepeatModeler2 (v2.0.7) [28] and RepeatMasker (v4.1.9) [29]. Gene prediction was conducted in OmicsBox v3.4.5 (BioBam Bioinformatics, Valencia, Spain) using the Genome Analysis module with *Arabidopsis thaliana* selected as the closest annotated organism to guide *ab initio* predictions. As external protein evidence, we incorporated a curated set of Heliantheae protein sequences downloaded from NCBI, reflecting the closest available high-quality proteomes within the Asteraceae. These datasets were used by OmicsBox to refine intron–exon boundaries and improve gene model support in the highly repetitive *S. gordilloi* genome. The resulting structural annotation was exported as a GFF3 file and included predicted genes, transcripts, exons, and CDS features.

Functional annotation of the predicted protein-coding sequences was carried out in OmicsBox using the InterProScan [30] pipeline with default parameters. InterProScan integrated domain assignments from Pfam, SUPERFAMILY, Gene3D, CDD, PRINTS, and PROSITE, and retrieved Gene Ontology terms, pathway associations, and InterPro entries for each protein model. To improve the biological accuracy of the annotation and reduce inflation arising from fragmented or transposable-element–derived predictions, we applied a two-step post-processing filter using AGAT v1.6.1 [31]. First, we removed all gene models whose total CDS length was <300 bp to eliminate truncated ORFs and spurious short predictions. Second, we parsed the InterProScan TSV output to identify entries associated with transposable element activity (for example transposases, integrases, retrotransposon-related domains) and removed the corresponding transcript and gene models from the GFF3. This filtering resulted in a substantial reduction of low-confidence predictions and yielded a final high-confidence gene set appropriate for downstream evolutionary and comparative analyses.

## 7. Data Availability

The raw sequencing reads generated in this study are available in the NCBI Sequence Read Archive (SRA) under accession number SRR30429356, within BioProject PRJNA1152989.

The genome assembly has been deposited in NCBI under BioProject PRJNA1152989. The genome annotation files generated in this study are available in GigaDB.

### 8. List of abbreviations

bp: Base pairs
BUSCO: Benchmarking Universal Single-Copy Orthologs
CDS: Coding Sequence
Gb: Gigabases
kb: Kilobases
LTR: Long Terminal Repeat
Mb: Megabases
ONT: Oxford Nanopore Technologies
SRA: Sequence Read Archive
TE: Transposable Element

## 9. Declarations

## Ethics Approval

All sampling procedures were conducted under Research Permit No. MAAE-DBI-CM-2021- 0198 issued by the Ecuadorian Ministry of Environment. No endangered individuals were harmed during sampling, and all collections were performed in accordance with national regulations for research on native plant species.

## AI Use Statement

Generative artificial intelligence (AI) tools were used during manuscript preparation to assist with language refinement, grammar correction, and improvement of readability. No AI tools were used for data generation, analysis, interpretation, or drawing scientific conclusions. All outputs generated with AI assistance were carefully reviewed and edited by the authors, who take full responsibility for the accuracy and integrity of the final manuscript.

## Authors’ contribution

GP: Conceptualization, Formal analysis, Funding acquisition, Investigation, Methodology, Writing – original draft, Writing – review & editing.

GRT: Conceptualization, Funding acquisition, Investigation (field sampling), Writing – original draft, Writing – review & editing.

EVD: Investigation, Formal analysis, Writing – original draft.

DB: Investigation, Formal analysis, Writing – original draft.

MLT: Conceptualization, Funding acquisition, Methodology, Supervision, Writing – review & editing.

## Acknowledgements

We thank the staff of the Galapagos Science Center for logistical support during fieldwork, especially Paul Yépez for his essential role in sample collection. We also thank Corbin Jones for his valuable guidance. We are grateful to members of the Plant Biotechnology Laboratory at Universidad San Francisco de Quito (USFQ) for their assistance and support throughout this project. Finally, we thank the Galápagos National Park Directorate and the Ecuadorian Ministry of Environment for granting research permits.

